# Neural dynamics of the impact of font style on lexical decision making in adult dyslexia

**DOI:** 10.1101/494724

**Authors:** Léon Franzen, Marios G. Philiastides

## Abstract

Good reading comprehension is indispensable in many situations including contract-based transactions that have become so prevalent in our everyday lives. People with dyslexia often exhibit impairments in this important cognitive process. Although the effects of italics – a commonly used style for highlighting important content in a range of documents – and font in general, have been explored with behavioural measures, their impact on human brain dynamics remains poorly understood. Here, we used electroencephalography (EEG) to investigate the specific effects of italics in a sentence reading lexical decision task in adult dyslexics and an age matched non-dyslexia group. Overall, the performance of dyslexics was worse. Cluster-based event-related potential (ERP) analysis revealed that brain responses within the first 300 ms following the decision stimulus differed in amplitude and spatial distribution between dyslexics and non-dyslexics when processing italicised text. An initial ERP component over occipitotemporal electrode sites started to differ between the groups as early as 167 ms following the onset of short italicised decision words. A subsequent ERP component over centrofrontal electrodes showed differences lasting until about 300 ms post-stimulus onset. Inter-individual amplitude differences in this centrofrontal neural signal were predictive of behavioural performance across participants, further highlighting the role of fast post-sensory linguistic processes in lexical decision making. Crucially, our findings emphasise the importance of choosing font style carefully to optimise word processing and reading comprehension by everyone.

## 1 Introduction

High levels of literacy are essential in many social and economic exchanges. This is particularly the case for exchanges involving legal contracts, which comprise almost exclusively large amounts of text. When dealing with these documents, impaired orthographic and semantic processing during reading can have detrimental consequences. One group that often falls short of developing efficient reading processes includes people diagnosed with dyslexia, a heterogeneous learning disability characterised by, inter alia, deficits in acquisition of efficient reading (Lyon et al., 2003).

These general behavioural deficits increase as readability decreases with certain types of fonts – also known as typefaces (Rello and Baeza-Yates, 2016, 2013). Italic fonts in particular play a crucial role in contracts, where they are commonly used to emphasise important content (Adams, 2013) as it is believed they help facilitate reading comprehension and retentiveness. Yet dyslexics exhibit a strong aversion and compromised reading performance with italic fonts (Rello and Baeza-Yates, 2016, 2013) compared to non-dyslexics. These deficits are exacerbated when dyslexics are required to read under time pressure – as is often the case with many legal documents – since they have difficulties with fast visual information processing (Warnke, 1999) and reading speed (Lefly and Pennington, 1991). As such, the use of italicised font style in particular is discouraged (British Dyslexia Association, 2018).

Previous human electroencephalography (EEG) studies suggest that the efficient visual processing of words is a fast and incremental neural process (Rayner and Clifton, 2009), with important orthographic and (sub)lexical steps being performed within 250 ms after encountering a word (Dien, 2009). Correspondingly, differences between meaningless false font strings and actual words (Proverbio et al., 2004) as well as differences associated with changes in font type (Chauncey et al., 2008) have been reported as early as 150 ms post-stimulus in non-dyslexics.

Dyslexics’ difficulties with fast orthographic word processing have also been linked to differences in early temporal components such as the P150/P1 (Araújo et al., 2012; Taroyan and Nicolson, 2009), the word N170/N1 in children (Hasko et al., 2013; Maurer et al., 2011, 2007) and adults (Brem et al., 2006; Helenius et al., 1999, 1998; Maurer et al., 2008; Salmelin et al., 1996; Schlaggar and McCandliss, 2007). More specifically, the word N170/N1 component has been associated with the so-called ‘visual word form area’ (e.g., Brem et al., 2006), located in the left posterior regions of the fine grained reading network (Price and Devlin, 2011; Price and Mechelli, 2005; Shaywitz et al., 2003), and is believed to play an important role in the fast visual processing of word shapes (McCandliss et al., 2003; Schlaggar and McCandliss, 2007).

While these findings demonstrate the presence of early neural correlates of visual word processing, the temporal dynamics underlying changes in font style in complex reading tasks along with their effects on lexical decision making between dyslexics and non-dyslexics remain poorly understood. Here, we collected EEG data during a sentence reading lexical decision task – presented in the context of legal language – to investigate differences in the brain dynamics of adults with and without dyslexia during processing of italic font and to demonstrate how these lead to changes in behaviour.

## 2 Materials and Methods

### 2.1 Participants

Fifty-one (28 dyslexics, 23 controls) male, right-handed, native English-speaking adults participated in this study (*Mean age*_*dyslexics*_ = 22.68, *SD*_*dyslexics*_ = 4.10 and *Mean age*_*controls*_ = 24.22, *SD*_*controls*_ = 5.84). Twenty-eight of those were diagnosed with dyslexia as identified by providing proof of an official diagnosis by a qualified specialist. The age they were given their diagnosis ranged from 5 to 30 years. All subjects had self-reported normal or corrected-to-normal vision, and reported no history of neurological disorders. Participants were current or former university students. Written informed consent was obtained from all participants in accordance with the guidelines of the Centre for Cognitive Neuroimaging at the University of Glasgow. All participants were paid £12 for their participation. This study was approved by the ethics committee of the College of Science and Engineering at the University of Glasgow (ethics application CSE300150102).

Only right-handed adult participants were recruited due to differences in prevalence of left-hemisphere language dominance based on handedness (Pujol et al., 1999). This was of particular importance since recent findings show differences in functional brain organisation and lateralisation in adult dyslexics (Finn et al., 2014). Additionally, we included only male participants due to the prevalence of dyslexia in men (gender ratio of 4:1) (Shaywitz et al., 1990). From the original group, four participants had to be excluded from the analysis. One control participant was excluded due to an excessive number of no-choice trials (i.e., 50 trials). The other three participants, two controls and one dyslexic, were excluded due to excessive noise in the recorded EEG data.

We administered a questionnaire to all participants with dyslexia upon finishing the task, which revealed that the majority of the dyslexics (78%) did not consciously perceive the difference in font style (i.e., Arial regular vs italic) during the experiment at all, or only did so towards the very end of the experiment. In contrast, all but one of the control participants indicated that they had recognised the difference in font style. This information was used for defining subgroups during subsequent analyses of the behavioural and neural data. This subdivision resulted in three groups with 21 dyslexic non-recogniser (DYS NO-R.), 6 dyslexic recogniser (DYS R.) and 20 control (CON) participants. Importantly, we used the comparison of the dyslexic non-recogniser to the control group as our main group contrast. Due to its small sample size, we treated the dyslexic recogniser group separately mainly for reference purposes only and to keep the main dyslexic non-recogniser group as homogenous as possible.

### 2.2 Stimuli and experimental procedure

Our task combines the comprehension of meaningful sentences as in Connolly et al. (1995), with a final decision on one congruent or incongruent real word (i.e., the ‘decision word’), made after the sentence itself. Entire sentences taken from various real-life legal contracts with differing content were presented using the Rapid Serial Visual Presentation (RSVP) technique (e.g., Rayner and Clifton, 2009). Specifically, participants were shown each sentence, centrally, word-by-word at a speed of 200 ms per word (Fig. 1a). This speed corresponds to the average reading speed of a skilled reader reading between 250 and 350 words per minute, and approximates the average fixation duration when reading text (Rayner and Clifton, 2009; Sereno and Rayner, 2003). Overall, participants were presented with 20 practice and 320 experimental sentences of twenty words and one decision word each. Note that only experimental sentence trials were analysed. Although the RSVP technique does not allow for a preview benefit, it was used in order to reduce the likelihood of introducing confounding eye movement artefacts into the EEG data, which are typically observed during regular sentence reading (Eden et al., 1994; Prado et al., 2007).

**Fig. 1.**
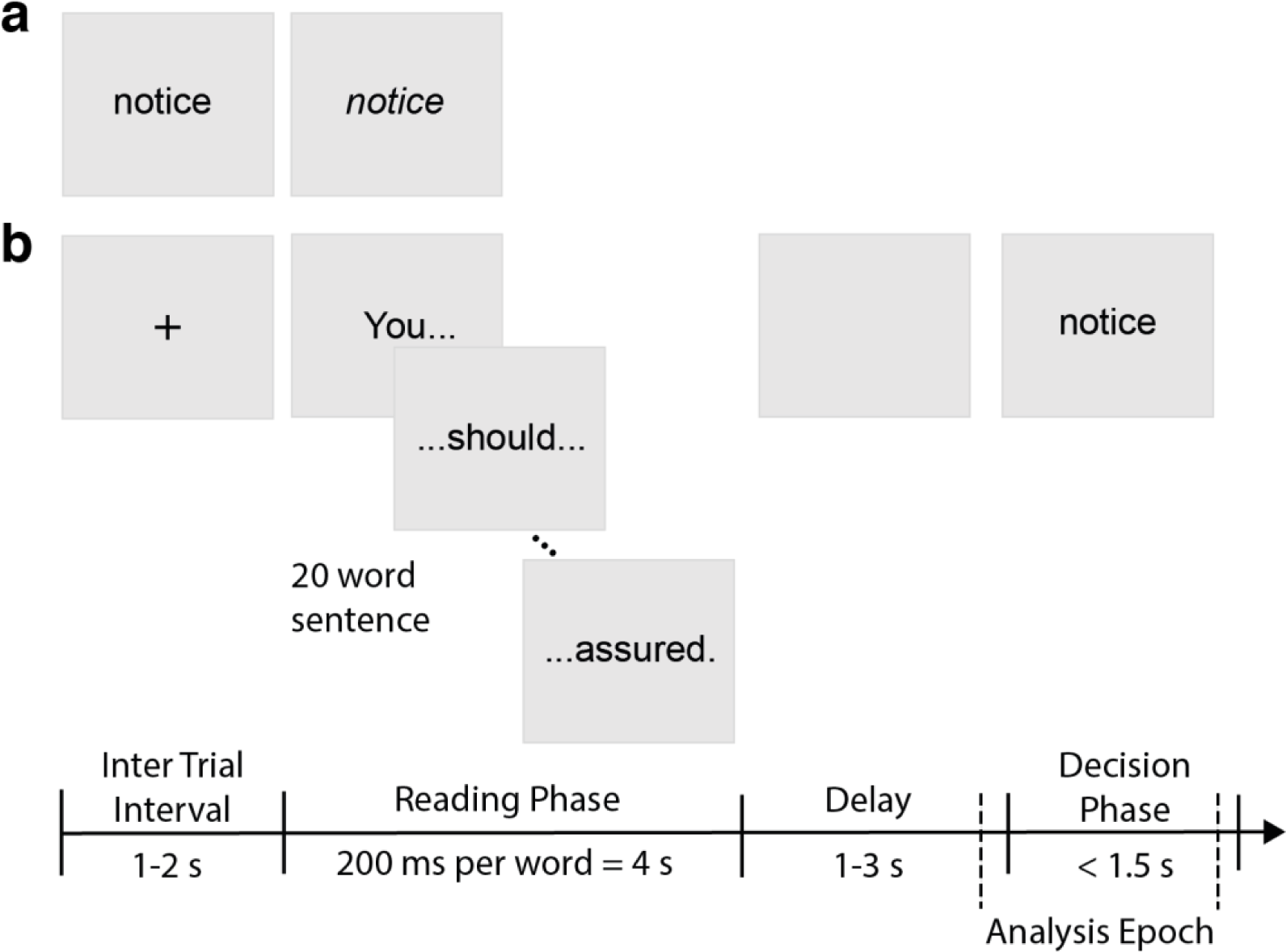
Experimental design and example of stimuli. a) Left: example of word in Arial regular font; right: example of word in Arial italic font. All words were presented as shown in black Arial font on a light grey background. b) Schematic representation of the experimental task showing the order of presented events for one trial. Participants were presented with a sentence of 20 words using the RSVP technique (200 ms per word) during the ‘Reading Phase’ and had to decide whether the decision word in the ‘Decision Phase’ was presented in its preceding sentence. A blank grey screen of variable delay (1-3 s) was shown in between the offset of the sentence and the onset of the decision word. Participants had up to 1.5 s to indicate their choice during the ‘Decision Phase’ starting with the onset of the decision word. Dashed vertical lines indicate the time period of the epoch included in all subsequent EEG analyses. Each trial was followed by an ‘Inter Trial Interval’ that varied randomly between 1 and 2 s.

Following the presentation of each sentence and after a short, jittered delay period (between 1 and 3 seconds), a decision word was presented for 1.5 seconds. Participants were instructed to indicate whether the decision word was included or not in the preceding sentence by pressing one of two keys with their index or middle finger of their right hand, respectively. The decision words were counterbalanced for high and low word-frequency between font styles (see Table 1 for examples of word-frequency) as assessed by the British National Corpus (BNC) frequency per million words (version 3; BNC Consortium, 2007). A written word-frequency of <40 per million was taken as cut off value for low-frequency words. In addition, their character length, ranging between 2 and 13 letters, was balanced between conditions in order to exclude confounding effects of word length (see Table 1 for examples of word length; Hauk et al., 2006; Paizi et al., 2013). Only words appearing between position 5 and 15, within their respective 20-word sentence, qualified as decision words. The delay time between encountering the decision word within the sentence – if included – and its appearance as the decision word was kept constant at a rate of 4 seconds. The decision word was either included in the preceding sentence or presented in a different sentence throughout the experiment. However, we ensured that decision words not included in their preceding sentence were matched with the sentence on a contextual level as well as possible (see Table 1 for examples of congruency). A trial was counted as missed and excluded from the analysis if no response was given within the available 1.5 second decision period. Across all participants and trials, this resulted in 2% of trials being excluded.

**Table 1.** Example of relevant stimuli conditions. Sentences were presented word-by-word at a rate of 200 ms per word. The decision word was shown for a maximum of 1.5 seconds.

| Condition | Sentence | Decision Word |
| --- | --- | --- |
| Short word of high-frequency (congruent) | For the Annual Medical Plan, the first travel date must fall within three months of purchase of the corresponding insurance. | fall |
| Short word of low-frequency (incongruent) | *However, C2Phone will not accept any liability for items damaged in transit, therefore the usage of recorded postage is recommended.* | repair |
| Long word of high-frequency (congruent) | *On receiving an application, the committee will consider if it is appropriate to determine a market rent as a reference.* | committee |
| Long word of low-frequency (incongruent) | The customer must take adequate steps to look after the leased equipment appropriately and maintain it in a satisfactory condition. | precautions |

All sentences and their subsequent decision words were shown in black writing in one of two distinct font styles – Arial regular or Arial italic – on a light grey background (RGB value: [128, 128, 128]). This change in Arial font style served as a perceptual manipulation. We chose Arial font, as it is a sans-serif font that is frequently used in contracts. In addition, Arial regular font has been used in previous behavioural and eye-tracking experiments, and appears to be easily legible for dyslexics (Rello and Baeza-Yates, 2016, 2013). Thus, the task should be less challenging for dyslexic participants, and put emphasis on the italic font style manipulation without adding serifs as a confounding factor.

The experiment was designed and run using PsychoPy (version 1.83; Peirce, 2008). The 320 experimental trials were split into four blocks that each contained 80 sentences each. Both conditions – Arial regular and Arial italic – were counterbalanced and sentences randomised within each block. Upon finishing the task, dyslexic participants were asked to complete a questionnaire with items referring to their dyslexia including age of diagnosis, past and current symptoms as well as task inherent properties such as the conscious recognition of italics during the experiment.

### 2.3 Behavioural analysis

For our behavioural analysis, we split decision words into two bins based on their word length. Accordingly, we henceforth refer to ‘short words’ (≤ 6 letters) and ‘long words’ (> 6 letters). This distinction is important because of reports showing evidence for the perception and processing of short words of up to 6 letters as one unit (McCandliss et al., 2003). Since we abstract letter identities to identify words (Sanocki and Dyson, 2012), it is likely that distinct font styles affect a word’s orthographic percept differently dependent on its length by altering its shape or word form (i.e., its contour line). Changing the font style from regular to italic alters a word’s perceived shape only slightly by manipulating its angle whilst keeping most other font properties constant. This change seems to be particularly relevant for short words that can be quickly decoded as one unit with a single fixation whilst having relatively similar phonologic properties. Longer words often stand out perceptually merely due to their length whereby the effect of a small change in font style, as used in this study, might become secondary or even get abolished.

We quantified behavioural effects using separate Generalised Linear Mixed Effects Models (GLMM) for decision accuracy and response time using the *lme4* package (Bates et al., 2015) in *RStudio* (RStudioTeam, 2016), and specifying a *binomial logit* and a *gamma* model, respectively. These models used the maximal random effects structure justified by the design including random correlations (Barr et al., 2013). Furthermore, both models included all main effects and interactions of the three predictors: *group* (control, dyslexic recogniser and dyslexic non-recogniser), *font style* (Arial regular and Arial italic) and *word length* (short and long) as well as by-subject and by-item random slopes and random intercepts for all relevant main effects and interaction terms. The three predictors were entered in mean-centred form. Given that the variable *group* had three levels, two different coding variables were required, treating the control group as a baseline group. Post-hoc likelihood-ratio *X*^2^ model comparisons were employed to evaluate the significance of the main effects revealed by the GLMM analysing their predictive power on decision accuracy.

### 2.4 EEG data acquisition

Continuous EEG data was acquired in an electrostatically shielded and sound attenuated room from a 64-channel EEG amplifier system (BrainAmps MR-Plus, Brain Products, Germany), with Ag/AgCl scalp electrodes placed according to the international 10-20 system on an EasyCap (Brain Products GmbH, Germany). In addition, all channels were referenced to the left mastoid during recordings and a chin electrode acted as ground. Input Impedance of all channels was adjusted to <50kΩ. Data was sampled at a rate of 1000 Hz and underwent online (hardware) filtering with a 0.0016–250 Hz analogue band-pass filter. Trial specific information including experimental event codes and button responses were recorded simultaneously with the EEG data using Brain Vision Recorder (BVR; version 1.10, Brain Products, Germany). These data were collected and stored for off-line analysis.

### 2.5 EEG data pre-processing

Off-line data pre-processing was performed with MATLAB 2015a (The MathWorks, 2015) by applying a software-based 0.5–40 Hz band-pass filter to remove slow DC drifts and higher frequencies (> 40Hz) as we were mainly interested in slower evoked responses that fall within the selected frequency range. These filters were applied non-casually (using MATLAB ‘*filtfilt*’) to avoid phase-related distortions. Additionally, the EEG data were re-referenced to the average voltage across all channels.

Subsequently, we removed eye movement artefacts such as blinks and saccades using data from an eye movement calibration task completed by participants before the main task. During this task participants were instructed to blink repeatedly upon appearance of a black fixation cross on light grey background in the centre of the screen before making several lateral and horizontal saccades according to the location of the fixation cross on the screen. Using principal component analysis, we identified linear EEG sensor weights associated with eye movement artefacts, which were then projected onto the broadband data from the main task and subtracted out (Parra et al., 2005). Trials with excessive noise in the EEG signal were rejected manually by visual inspection (< 3% of all analysed trials across participants).

### 2.6 Main EEG data analysis

To identify temporal activity related to orthographic word processing in the EEG data, we used a sliding window approach on our stimulus-locked ERP data. Namely, for every participant, data were averaged within time windows of 50 ms length, centred on specific time points across the epoch, starting at 100 ms prior to the presentation of the decision word and ending at 900 ms after its onset. These windows were shifted in increments of 3 ms. Our statistical analysis focused on stimulus-locked data lasting up to 600 ms post-stimulus onset, in order to avoid any period that might include confounding motor preparatory signals (*M*_*RT all trials*_ = 892 ms, *SEM*_*RT all trials*_ = 17 ms).

We tested for neural differences between the dyslexic non-recogniser and control groups employing a univariate non-parametric cluster-based permutation analysis as implemented in the LIMO-EEG-Toolbox (Pernet et al., 2011) for MATLAB. First, we averaged across trials separately for each within-subject condition (Arial regular and Arial italic) and each subject, to identify relevant EEG components independent of word length. Second, we proceeded to compare ERP amplitudes between the two groups of interest (dyslexic non-recogniser and control) to identify contiguous spatial and temporal clusters exhibiting differences in ERP amplitude using a non-parametric permutation analysis, while at the same time correcting for multiple comparisons (Maris and Oostenveld, 2007; Pernet et al., 2015).

Specifically, every sample from an electrode-time pair was compared between the two groups; separately for each font condition. Our permutation procedure involved randomly shuffling participants without replacement in order to generate two new random groups by reassigning each subject to one of the two testing groups. Sampling without replacement was chosen due to reports of this approach being most reliable for a limited number of samples (Pernet et al., 2015). We repeated this procedure 1000 times and performed between-group two-sample t-tests for every sample from each electrode-time pair. Samples exceeding a threshold of α < .05 were grouped in spatiotemporal clusters according to a neighbourhood matrix. This matrix specified spatial adjacency between electrodes using 3.7 cm as the maximum neighbourhood distance. We set the minimum number of significant channels in a cluster to 2. Next, a maximum cluster-mass permutation distribution was obtained by recording the sum of t-values of the maximum significant spatiotemporal cluster for each of the 1000 random permutation iterations. This resulted in a randomised cluster-level summary statistic under the null-hypothesis that was used to determine an appropriate threshold (p < .05) for assessing statistical significance of the differences in the original data.

This analysis was complemented by unbiased effect size calculations (Hedges’ g for between-subject designs as described in Lakens, 2013), which compared differences in mean amplitude for significant spatiotemporal clusters between dyslexic non-recognisers and controls (henceforth, *g*). The interpretation of Hedges’ g effect sizes is comparable to the benchmarks reported for Cohen’s *d* (0.2 = small effect, 0.5 = medium effect, ≥ 0.8 = large effect; Cohen, 1988). In this context, an effect size of 0.5 can be interpreted as a difference of two means by half a standard deviation (Lakens, 2013). Effect size calculations for one-sample tests were performed using the ‘*mes*’ function of the Measures of Effect Size Toolbox (Hentschke and Stüttgen, 2011) in MATLAB.

We repeated this procedure for trials displaying only short decision words – Arial regular short and Arial italic short – to identify specific perceptual neural components that are simply a result of perceptual effects of italic font during lexical decision making. Due to physiological acuity limitations, only high spatial frequency information of short words can be extracted in its entirety with a single fixation (Rayner, 1998). Letters of short words extracted during a fixation are processed in parallel and as such their visual word form gains importance as they are believed to be perceived as one unit (McCandliss et al., 2003). Short words are also psycholinguistically more consistent and thereby can make our perceptual effect of font more trackable. In contrast, long words are often perceptually more salient simply due to their length and rarer occurrence in written text. In their case, gaze duration increases almost linearly with the number of letters (Rayner et al., 1996). For these reasons, and to avoid diluting our perceptual effect of font by these factors, we also tested short words separately.

Furthermore, since trial-to-trial changes are of particular importance when attempting to link neural activity to visual stimulus parameters (Rousselet et al., 2008) and decision making (e.g., Philiastides and Sajda, 2007), we complemented our ERP analysis with a robust multilinear single-trial regression analysis (using MATLAB ‘*robustfit’*). This single-trial regression approach examined whether activity in neural components of interest (identified by means of our cluster-based permutation analysis) observed over distinct electrode and temporal clusters, covaried together across individual trials. This analysis computed (*β*) parameter estimates by regressing z-scored single-trial peak ERP amplitudes of separate spatiotemporal clusters against each other. These single-trial ERP peak amplitudes were computed by averaging ERPs across all electrodes within the relevant electrode clusters before selecting the peak amplitude (using MATLAB ‘*max*’ or ‘*min*’) within a 100 ms window around a component’s group grand average peak time. To quantify a potential functional link between our distinct spatiotemporal neural components, we compared the resulting β parameter estimates (across subjects) against the null-hypothesis (that is, β parameter estimates come from a distribution with mean equal to 0) using a two-sided paired t-test.

Lastly, we employed robust bend correlations (Pernet et al., 2013) to test the extent to which activity in our neural components of interest (using peak amplitudes computed from each participant’s ERP grand average) could predict behavioural mean accuracy on italic short word trials across all participants. We also used robust bend correlations for correlating individual mean response time with each component’s peak time (using the peak timing of the previously identified peaks) separately across participants. This part of our analysis intended to rule out differences in response time as the main reason for the observed neural differences.

## 3 Results

### 3.1 Behavioural results

Overall, mean accuracy was at least 67%, independent of group or condition illustrating all groups were able to perform the task well above chance (Fig. 2a). First, we ran a mixed effects model for accuracy with predictors: *group* (control, dyslexic recogniser and dyslexic non-recogniser), *font style* (Arial regular and Arial italic) and *word length* (short and long) and their interactions.

**Fig. 2.**
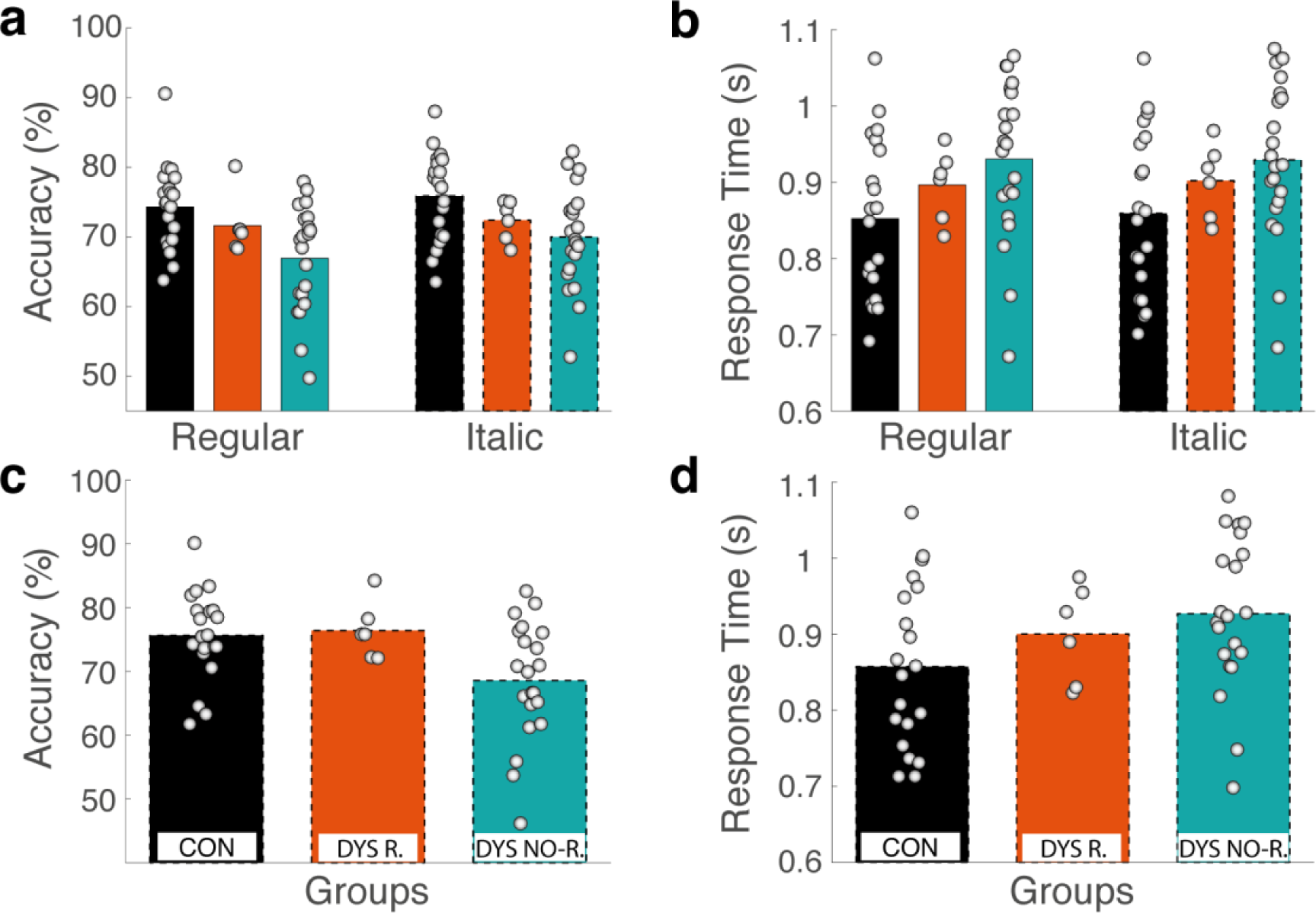
Behavioural performance and response time of all participants separated by group. Panels a and b illustrate group means for all trials collapsed across word length, whereas panels c and d present group means only for trials presenting short decision words in italic font. Dots denote each participant’s individual mean across trials in the respective condition. Black bars depict mean values for the control (CON), orange for the dyslexic recogniser (DYS R.), and green for the dyslexic non-recogniser (DYS NO-R.) group. a) Mean values for accuracy collapsed across word length. Bars surrounded by a solid outline represent trials in Arial regular font, a dashed outline represents Arial italic trials. b) Mean values for response time collapsed across word length for Arial regular and Arial italic trials. Bar outlines and colours as in panel a. c) Mean decision accuracy for short italicised decision words (≤ 6 letters). d) Mean response times for decisions on short italicised decision words.

We found main effects of *group* (*Z* = −3.43, *p* < 0.001; Table 2) and *font style* (*Z* = - 2.34, *p* = 0.0195; Table 2), without any significant effects of word length or interactions (for details, see Table 2; Fig. 2a). The group effect was specifically found for the contrast between the control and dyslexic non-recogniser group, with dyslexic non-recognisers showing significantly worse performance [*M*_CON_ = 75.11, *SEM*_CON_ = 0.55; *M*_NO-R_ = 68.45, *SEM*_NO-R_ = 0.59] across all conditions (Table 2; Fig. 2a and 2c). These two main effects were confirmed via likelihood-ratio *X*^2^ model comparisons, which separately contrasted the goodness of fit of a full model with all predictors against our alternative model without one of the two main effects. Removing the main effect of the dyslexic non-recogniser group or font style from the full model each decreased the goodness of fit significantly (*X*^2^ (1) = 10.5, *p* = 0.001 and *X*^2^ (1) = 5.38, *p* = 0.02, respectively).

**Table 2.**
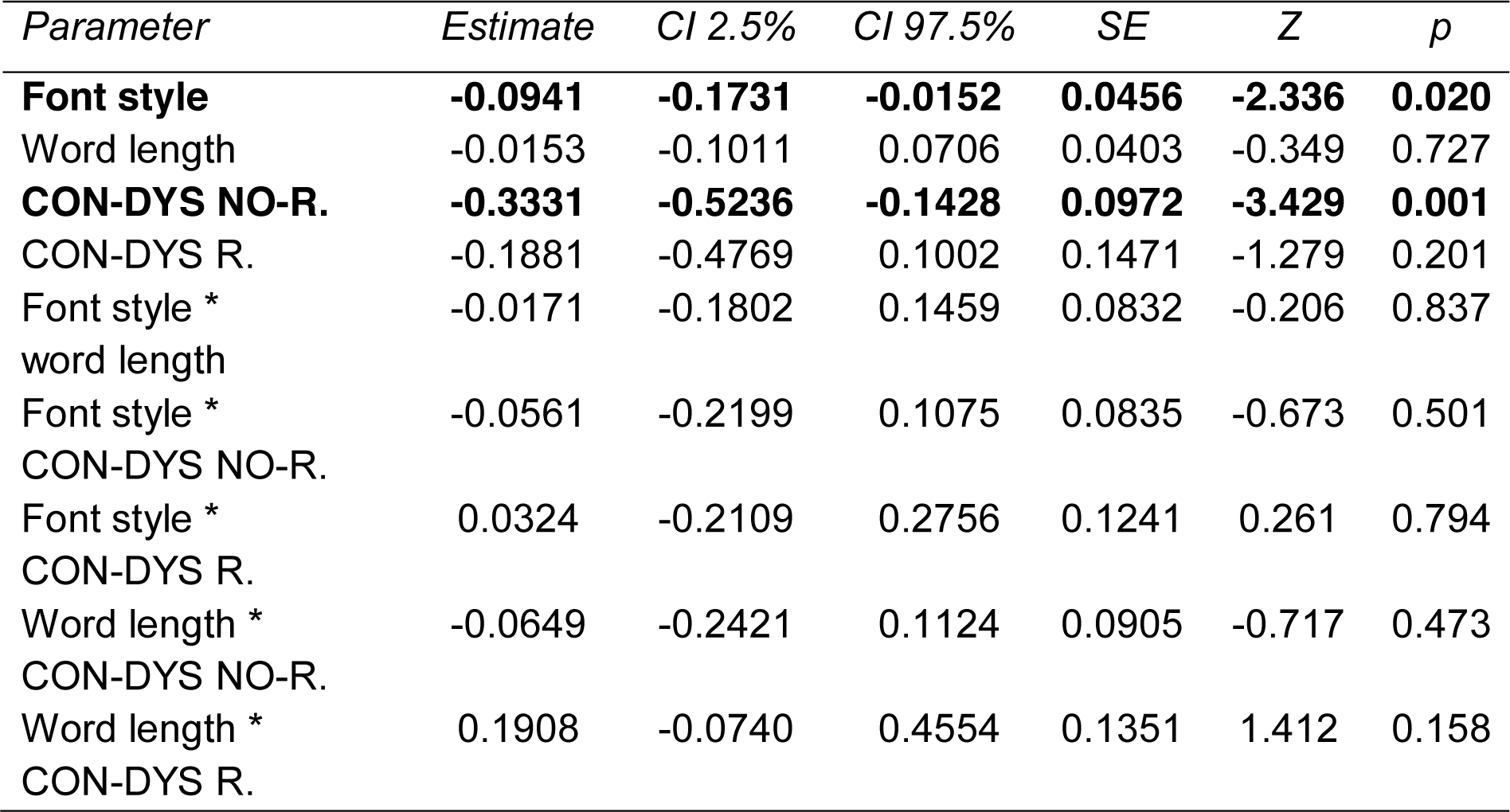
GLMM fixed effect parameter estimates for accuracy. Effects with significant predictive power after post-hoc likelihood-ratio *X*^2^ model comparisons in bold. Group labelling as follows: CON = controls; DYS NO-R. = dyslexic non-recognisers; DYS R. = dyslexic recognisers.

Next, we ran a mixed effects model for response time with the same three predictors and their interactions as the previous model but found no significant main effects or interactions (for details, see Table 3; Fig. 2b). Note, however, that on average there was a systematic trend for the dyslexic non-recognisers to indicate their decisions with longer response times (*t* = −1.76, *p* = 0.079; Table 3; Fig. 2b and 2d).

**Table 3.**
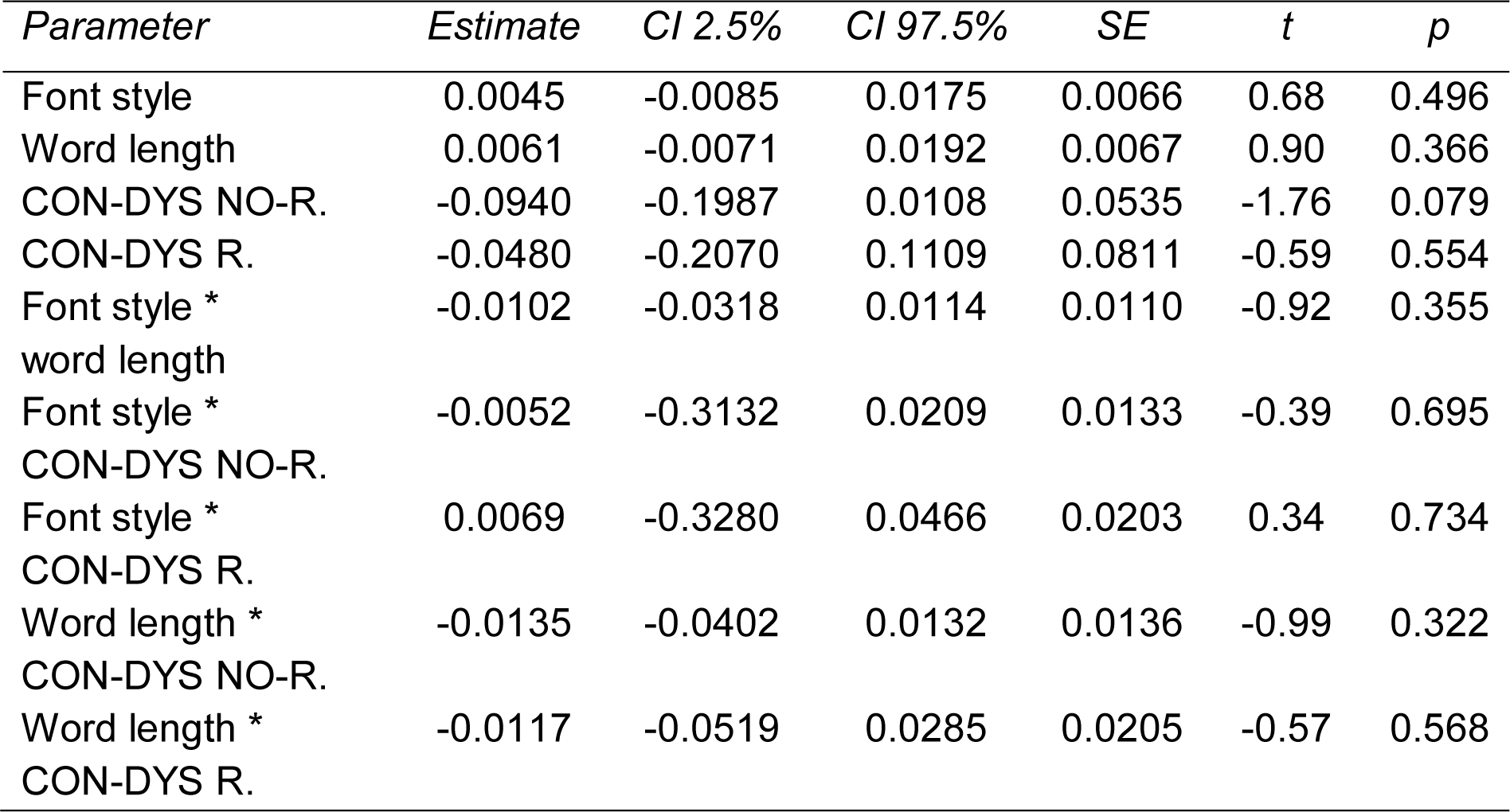
GLMM fixed effect parameter estimates for response time. Group labelling as follows: CON = controls; DYS NO-R. = dyslexic non-recognisers; DYS R. = dyslexic recognisers.

### 3.2 EEG results

We performed pairwise comparisons of the control and dyslexic non-recogniser groups of all italic font decision words to identify the general effect of this specific font style irrespective of word length. This analysis revealed significant differences between 215 and 281 ms post-stimulus onset (*M*_*diff*_ = 0.9 μV; 95% CI_MeanDiff_ [0.49, 1.53]; *g*_*italic words*_ = 1.01; Fig. 3a). The resulting ERP component differed significantly over right centrofrontal electrode sites (henceforth, centrofrontal component) as identified by our permutation analysis (Fig. 3b).

**Fig. 3.**
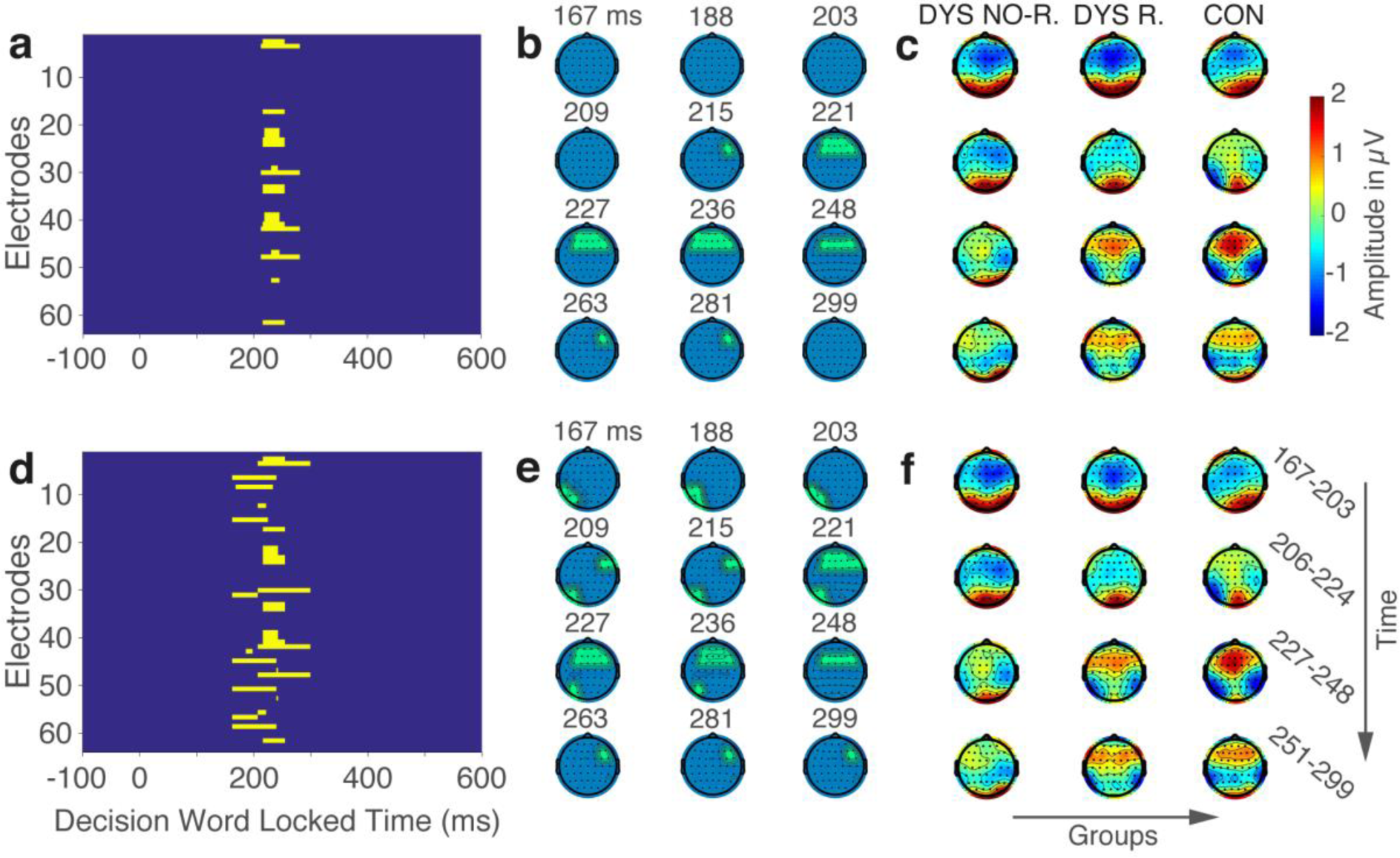
Temporal evolution and scalp distributions of ERP components. Panels a-c show ERP components for all italicised decision words independent of word length, whereas panels d-f illustrate the ERP components for trials presenting short decision words (≤ 6 letters) in italic font. Panels a, b, d and e depict significant ERP components for the main between-group contrast comparing the control to the dyslexic non-recogniser group on italic trials. a) Significant electrodes over time after cluster-based correction for multiple comparisons. Yellow indicates a significant electrode at a given time point. b) Evolution of significant ERP components over time and space at representative time points. Green denotes significant clusters of electrodes. c) Scalp distribution of activity averaged separately across participants of each group and time points corresponding to the same row in panel b and as specified in panel f. Groups are denoted as control (CON), dyslexic recogniser (DYS R.), and dyslexic non-recogniser (DYS NO-R.). d) Significant electrodes over time for italic short decision word trials after multiple comparisons correction. Yellow indicates a significant electrode at a given time point. e) Scalp distribution of significant electrodes at representative time points. Green denotes significant electrode clusters. f) Scalp distribution of activity averaged separately across participants of each group and time points corresponding to the same row in panel e. Group specification as in panel c.

To examine the precise perceptual effects of italic font without confounding effects of word length, we repeated the same between-group contrast but focused only on short italicised decision word trials instead (Fig. 3d-f). This contrast yielded significant differences in activity within an even wider time window extending from 167 to 299 ms post-stimulus onset (Fig. 3d). By focusing only on short italicised words, our permutation analysis uncovered an additional cluster of left occipitotemporal electrodes (henceforth, occipitotemporal component) whereby we observed a significant difference in ERP amplitude between dyslexic non-recognisers and controls earlier in time (i.e., 167 to 236 ms), compared to the later centrofrontal component (Fig. 3d and 3e). The difference between groups in this additional occipitotemporal component peaked at 175 ms post-stimulus (*M*_*diff*_ = 1.48 μV; 95% CI_MeanDiff_ [0.70, 2.33]; *g*_*occipitotemporal*_ = 1.09; Fig. 4a). We note for reference purposes that the profile of this occipitotemporal component (i.e., timing and peak amplitude) was similar between the two dyslexia subgroups (i.e., dyslexic non-recognisers and dyslexic recognisers) within the significant time window revealed by our permutation test (i.e., 167 to 236 ms; Fig. 4a).

**Fig. 4.**
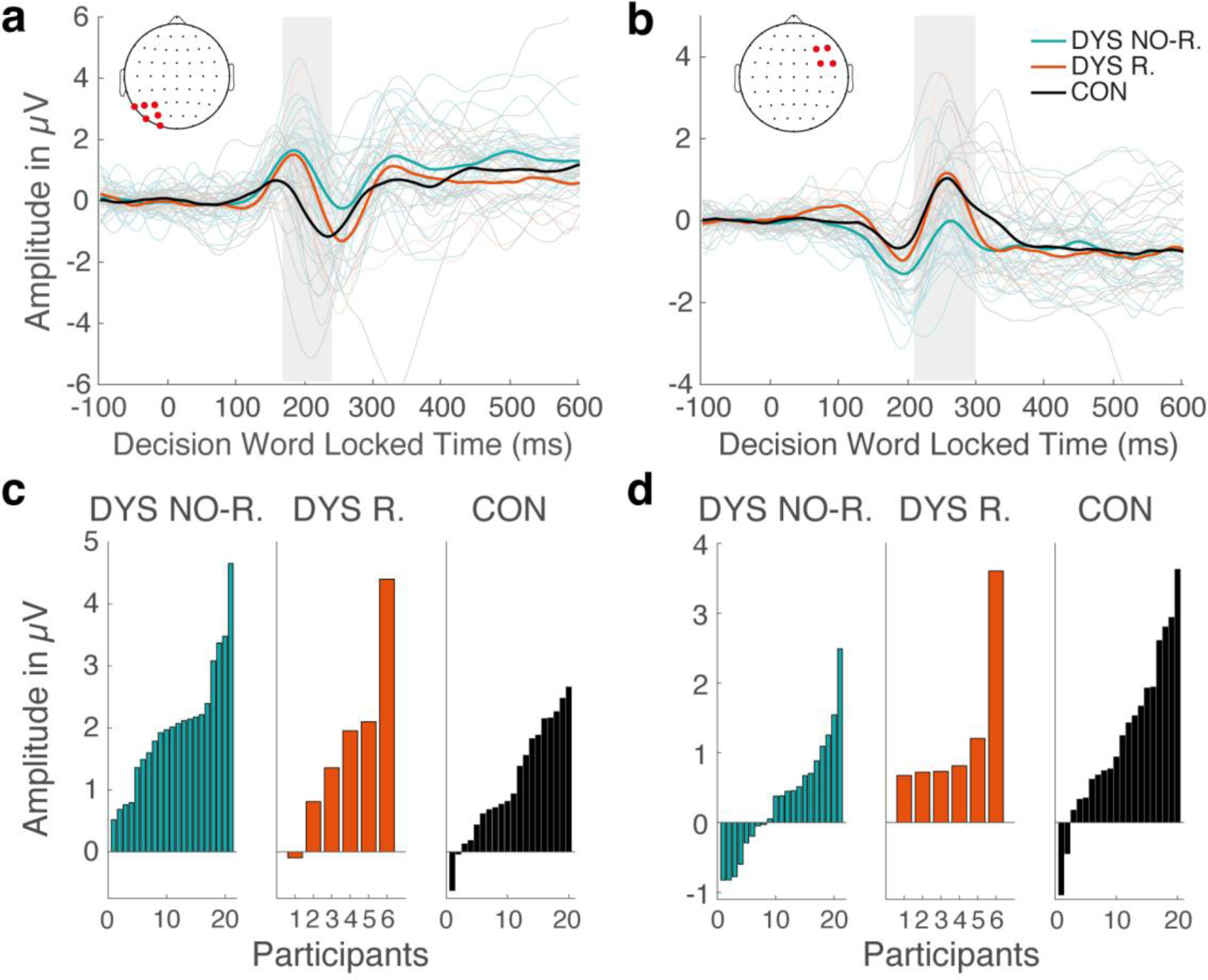
ERP group and individual participant grand averages and individual participants’ component peak amplitudes. a) Shows ERP group grand averages (thick lines) and individual participant grand averages (thin faded lines) averaged across electrodes of the significant left occipitotemporal cluster for short decision words in italic font. Averages for this cluster were computed across the following electrodes: P3, P5, P7, PO7, PO5, O1. Locations of these electrodes as shown in red on the corresponding scalp plot. b) Illustrates the same ERPs as panel a, but averages were computed across the four electrodes of the right frontal cluster that were consistent over time. Electrodes of this cluster were: F4, F6, FC4, FC6. Electrode locations of this cluster as shown in red on the corresponding scalp plot. c-d) Depict individual participants’ peak amplitudes sorted in ascending order by amplitude and separated by group and component (c, occipitotemporal component; d, centrofrontal component). Groups are denoted using the following colours in panel a-d: black for the control (CON), orange for the dyslexic recogniser (DYS R.), and green for the dyslexic non-recogniser (DYS NO-R.) group.

This analysis also captured the right centrofrontal component we identified in the original analysis with all word lengths. Figure 3e illustrates that this centrofrontal component started to differ significantly between groups at 209 ms post-stimulus, consistent with the original analysis. It is characterised by a difference in positive ERP peak amplitude between our main groups of interest around 250 ms post-stimulus over a cluster of right and central frontal electrodes (*M*_*diff*_ = 0.96 μV; 95% CI_MeanDiff_ [0.45, 1.50]; *g*_*centrofrontal*_ = 1.11; Fig. 4b). This component showed similar deflection and group grand average peak timing for all three groups (Fig. 4b). The two components overlapped between 209 and 236 ms post-stimulus onset but had different spatial topographies (Fig. 3e).

Figures 3c and 3f display the full scalp distributions of both ERP components. These differed clearly between the dyslexic non-recogniser and control groups throughout the time window associated with our two components. The control group showed lateralisation of activity depicted by opposite polarity over posterior electrodes early on, between 209 and 221 ms, as opposed to bilaterally distributed activity shown by both dyslexia subgroups on all italic trials (Fig. 3c). This lateralisation of activity started to emerge even slightly earlier than 209 ms for controls on short italic word trials (Fig. 3f). On these trials, any lateralisation of activity was absent in both dyslexia groups at all time points of the two components. Overall, the centrofrontal component (227 to 299 ms) in controls exhibited stronger positive activity over central anterior and stronger negative activity over right and left temporal electrodes as illustrated by their scalp maps. In contrast, dyslexic non-recognisers did not exhibit strong positive activation over anterior electrodes.

In contrast to the two components presented above that occurred within the first 300 ms after the onset of italicised decision words, we did not find any significant between-group ERP differences for time windows later than 300 ms post-stimulus or for between-group contrasts of trials presenting Arial regular font.

### 3.3 Single subject peak amplitudes

To avoid masking intra-group variance by comparing ERP group grand averages, we examined the individual subject peak amplitudes of the two previously identified components that fell into their respective time windows for short italic decision words. We found that participants’ peak amplitudes pointed consistently in the direction of their group’s grand average, independent of the component (Fig. 4c and 4d). Although, dyslexic non-recognisers showed slightly larger variability in polarity of their peak amplitudes of the centrofrontal component, the majority of single subject peak amplitudes pointed in the direction of their group grand averages (Fig. 4d). The consistency of the single subject peak amplitude polarity underlines that the observed group grand averages did not mask large variability within a group.

### 3.4 Linking activity between posterior and anterior neural components

Since there was a partial temporal overlap between our two identified components, we complemented our ERP analysis with a single-trial regression to evaluate a potential link between the sources underlying the relevant occipitotemporal and centrofrontal components (Fig. 3e). For every participant, we obtained standardised regression coefficients (βs) from a single-trial regression predicting peak amplitude of the centrofrontal component based on the peak amplitude of the preceding occipitotemporal component (Fig. 5a). We searched for a systematic correlation between our two component amplitudes by testing whether these β regression coefficients came from a distribution with mean zero using a two-tailed t-test. This procedure resulted in negative β coefficients for all participants independent of their group illustrating significant trial-by-trial functional coupling between the two identified neural components (*t*_46_ = −23.82, *p* < .0001; 95% CI [−0.4664, −0.3937]; *g*_link_ = 3.47; Fig. 5b). The negative direction of this relationship represents lower peak amplitude of the occipitotemporal component being coupled with higher peak amplitude of the centrofrontal component on a single-trial basis. Thus, this link illustrates that the identified components might form a part of an interconnected cascade of neural processes relevant for lexical decisions on short italicised words in particular.

**Fig. 5.**
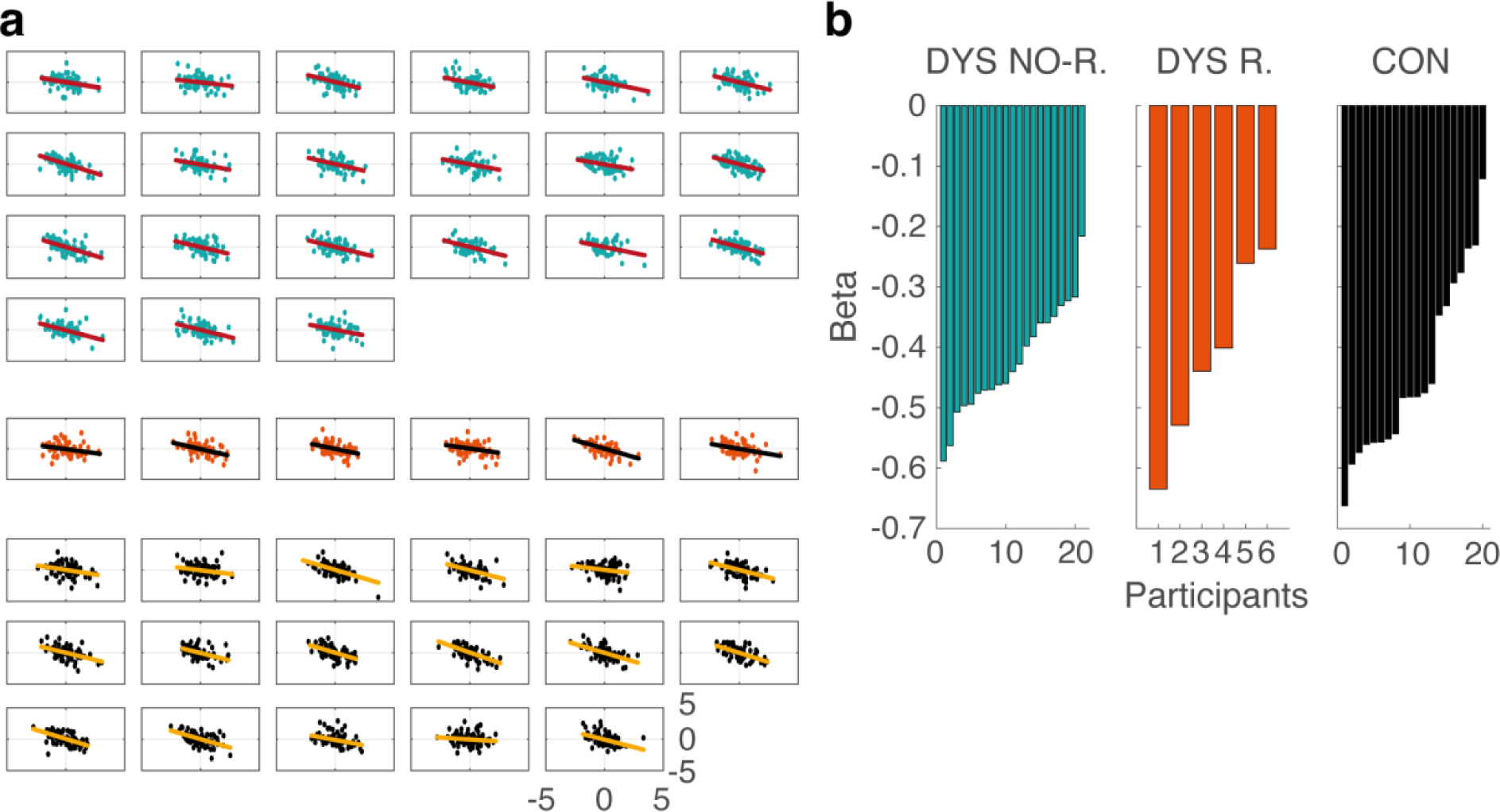
Single-trial peak amplitude regression. Results depicting the relationship between the left occipitotemporal and the right centrofrontal component. a) Single-trial peak amplitudes for the occipitotemporal and centrofrontal component by participant. One dot per trial. Coloured lines represent least-square lines. Groups are denoted by coloured dots and separated by space (from top: dyslexic non-recognisers (DYS NO-R.) in green, dyslexic recognisers (DYS R.) in orange and controls (CON) in black). b) Individual participants’ β coefficients separated by group and sorted by magnitude. Group colours as in panel a.

### 3.5 Peak amplitudes and decision accuracy

To test the extent to which the two identified ERP components were further predictive of behavioural performance, we correlated each of the two component peak amplitudes with mean decision accuracy across all participants. We found an interesting dissociation in the way the two components were linked to performance. Specifically, we did not find a significant relationship between the occipitotemporal component’s individual grand average peak amplitudes and mean accuracy across participants (*r*_45_ = -.16; *p* = .28; Fig. 6a). In contrast, subjects who exhibited higher centrofrontal mean peak amplitude also showed better behavioural performance independent of their group (*r*_45_ = .33; *p* = .024; Fig. 6b). Hence, the centrofrontal component represents one of the earliest processes during lexical decision making relevant for behavioural performance.

**Fig. 6.**
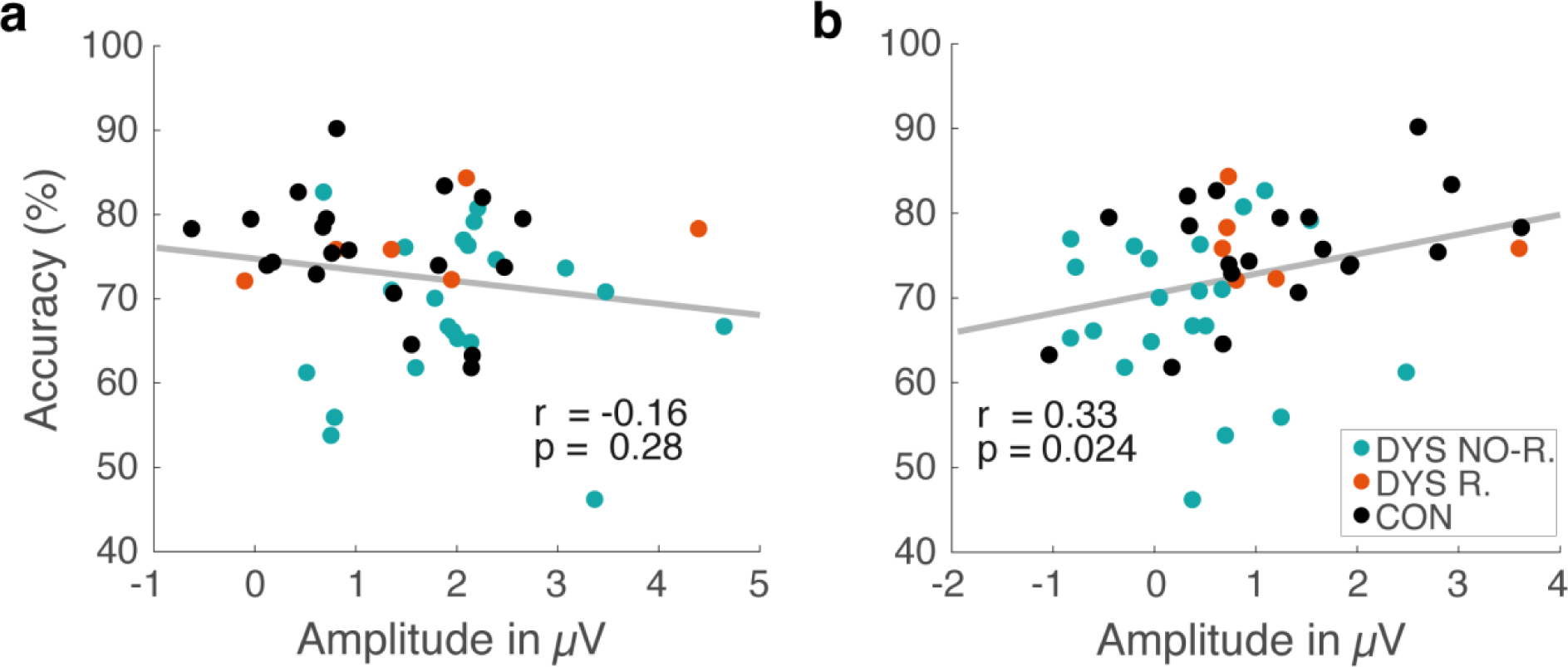
Correlation between individual participant mean peak amplitude and decision accuracy across all participants for short italic words separated by component. Colours of dots denote group affiliation. Groups: dyslexic non-recognisers in green, dyslexic recognisers in orange and controls in black. Statistical results from robust bend correlations using 20% bending in both directions (i.e., X and Y) across all participants are denoted next to the grey least-squares fit line. a) Occipitotemporal component peak amplitudes, computed in the window between 151 and 251 ms post-stimulus, correlated with mean decision accuracy. b) Centrofontal component peak amplitudes, computed in the window between 200 and 300 ms post-stimulus, correlated with mean decision accuracy.

### 3.6 Peak amplitudes and response time

Dyslexics are well known for responding slower on speeded decision tasks than controls due to processing speed deficits (Breznitz and Misra, 2003; McLean et al., 2011). Therefore, we intended to rule out the possibility that our identified neural components merely represented differences in response times between the two groups. We sorted all subjects by mean response time irrespective of group and plotted their individual ERP grand average activity for each of the two components separately. Both ERP component peaks were clearly observable locked to the stimulus and appearing within a 100 ms interval during and around the respective component’s time window, rather than shifting in time in accordance with longer response times (Fig. 7a and 7b). We quantified this observation using two robust correlations that evaluated the relationship between the participants’ mean response time and their occipitotemporal and centrofrontal component peak times separately across all subjects. Neither the occipitotemporal component (*r*_45_ = .13; *p* = .37) nor the centrofrontal component (*r*_45_ = .06; *p* = .70) were correlated with response time across participants. Consequently, the observed early neural components could not simply be explained by longer response times of the dyslexic non-recognisers. This account further endorses the proposition that the ERPs of the dyslexic non-recognisers and controls differed mainly in component amplitude.

**Fig. 7.**
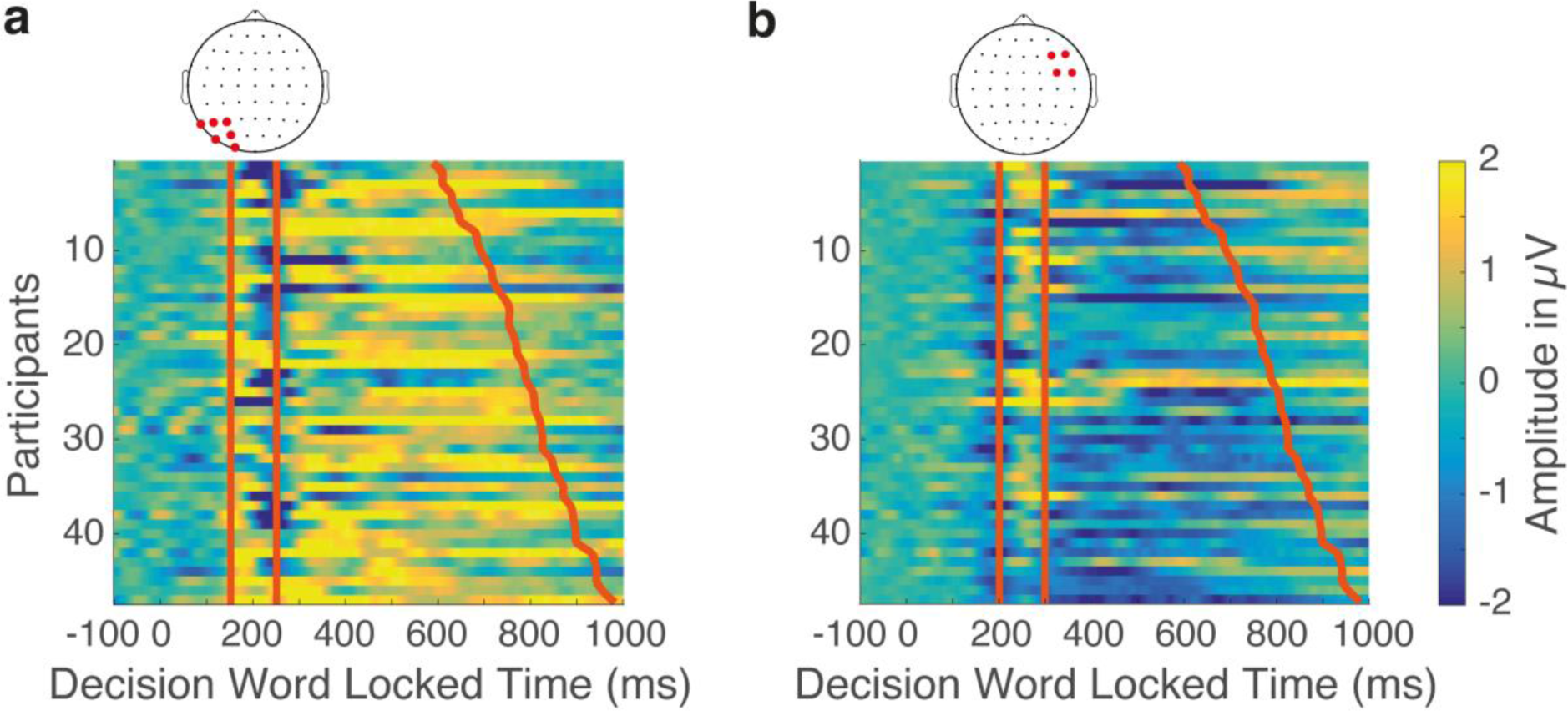
Individual participants’ mean ERP amplitude in relationship to their mean response time on italic short decision word trials. Vertical lines indicate the start and end of the 100 ms time window used for identifying peak amplitudes. Windows were 151 to 251 ms and 200 to 300 ms post-stimulus onset. Slanted orange lines depict mean response time. All participants were sorted in ascending order by response time independently of their group affiliation. a) Individual participant mean ERP amplitude averaged across the left occipitotemporal component’s electrode cluster shown in red on the scalp plot above. b) Individual participant mean ERP amplitude averaged across the temporally most consistent right centrofrontal electrodes shown in red on the scalp plot above.

## 4 Discussion

In this study, we have provided evidence for adult dyslexics’ impairments during a legal lexical decision making task when the text is presented in italic font. We linked behavioural impairments of dyslexics to two ERP components occurring within 300 ms post-stimulus onset that differed in amplitude between our groups. Crucially, we found a functional coupling between both components but only the second (later) centrofrontal component was more tightly linked to trial-by-trial changes in behavioural performance. Neither component shifted in time with participants’ response times. These two ERP components illustrate the challenges that italic font poses for adult dyslexics’ neural word analysis. The earlier occipitotemporal component substantiates the account that font affects the orthographic processing stages of lexical decision making, while changes in the later centrofrontal component are likely to reflect impairments in post-sensory processing more closely linked to the eventual decision outcome. Our results highlight the crucial role that font style plays within the word recognition cascade.

### 4.1 Behavioural impairments

Dyslexics demonstrated worse behavioural performance across font styles. In light of their well-known general deficits with text comprehension (e.g., Elbro and Petersen, 2004), we show here that these impairments persist into adulthood despite decades of reading practice. In our task, decision accuracy was higher for italicised words irrespective of group or word length. This finding is in line with reports of harder to read fonts’ – such as italics or Monotype Corsiva – ability to facilitate retention in non-dyslexics (Diemand-Yauman et al., 2010) and dyslexics (French et al., 2013). However, it stands in opposition to multiple reports of strong aversion against and worse performance on italicised font by dyslexics (Rello and Baeza-Yates, 2016, 2013). One plausible explanation for our finding is that overall participants might have allocated more attention (i.e., exogenous alerting or endogenous executive attention; Amso and Scerif, 2015) to the trials presenting italicised words as a consequence of their unfamiliar look, which could lead to increased salience in these trials. Salience is believed to be one of the key gatekeepers for attention allocation and bottom-up processing (Knudsen, 2007). In addition, the short time of each trial (∼4 seconds), leading to temporarily limited task demands, might have allowed even dyslexics to compensate in part by allocating more attention during more salient trials. It seems that encountering disfluent fonts in small chunks as in our and French and colleagues’ (2013) study does not necessarily pose a major problem for adolescent and adult dyslexics on the behavioural level. We argue that such improvements are most likely specific to tasks that only present short segments of words at a time and our specific sample of dyslexic university students. Based on dyslexics’ reported aversion against disfluent fonts, we can only speculate that the demands, and in turn (cognitive or ‘neural’) costs, would rise with increasing length of text passages in italicised or disfluent fonts.

Interestingly, the fact that only a small subset of dyslexic participants (22%) reported having recognised italic font during the experiment provides further evidence for dyslexics’ deficits with fast visual word recognition. However, the small size of this group, and the concomitant lack of statistical power, only allowed us to use this group for visualisation and reference purposes (relative to the other groups). A larger sample size would have allowed us to investigate intra-group differences of the spectrum disorder dyslexia in detail, particularly with respect to conscious font style recognition.

### 4.2 Changes in neural brain dynamics

#### 4.2.1 Occipitotemporal component

Our neural results suggest that dyslexics process italicised text differently. The earliest differential ERP component we found started at 167 ms post-stimulus onset for trials presenting short italicised decision words. This component was located over a cluster of occipitotemporal electrodes. Its timing strongly suggests that it represents different early sensory processing of italicised orthographic word forms, particularly, since we only observed it when changes in font style itself exert maximum impact on a word’s general shape (i.e., on short words). Visual word form perception is one of the first processes of fast and incremental visual word recognition in reading (Gaskell, 2007) whose orthographic analysis plays a crucial role in the word-recognition process (Lété and Pynte, 2003). In this respect, the efficiency and processing speed of perceiving the abstract letter and word identity is crucial for successful word recognition.

Importantly, effects of orthographic manipulations including words versus symbols (Appelbaum et al., 2009), transposed letters within a word (Grainger et al., 2006), different case (Spironelli and Angrilli, 2007), and font type (Chauncey et al., 2008) have been reported for an early latency range around 150 ms post-stimulus that is in line with our occipitotemporal component (Dien, 2009). Moreover, differential activity in the ventral stream, which hierarchically codes for letter strings (Vinckier et al., 2007), has been linked to deficits in two processes in dyslexia – neural adaptation (Perrachione et al., 2016) and efficient tuning to print (e.g., Kronschnabel et al., 2013; Mahé et al., 2012). These two processes affect early components within 250 ms and normally evolve during childhood in non-dyslexics (Brem et al., 2010). In combination with higher ERP amplitude being a signal for increased processing demands (Otten and Rugg, 2005), dyslexics’ higher occipitotemporal ERP amplitude suggests worse neural tuning and adaptation to print that persists into adulthood. Such sensory inefficiency may be a possible mediator of dyslexics’ word recognition impairments.

More support for this component’s role in orthographic word form perception comes from studies linking activity in the visual word form area, an area in left occipitotemporal cortex that plays a crucial role in word form recognition (e.g., McCandliss et al., 2003), to electrophysiological equivalents peaking around 170 ms (e.g., Brem et al., 2006). The visual word form area commonly shows aberrant blood-oxygen-level dependent (BOLD) activation patterns in dyslexia (Kronbichler et al., 2006; Shaywitz et al., 2003; Shaywitz and Shaywitz, 2005) and different BOLD activity for unfamiliar words (Wimmer et al., 2016). Our component’s occipitotemporal spatial distribution and its timing are consistent with these equivalents, however, methods with better spatial resolution such as fMRI are needed to confirm this interpretation.

Taken together, these observations suggest that (1) dyslexics’ exhibit deficits within the initial stages of orthographic processing, (2) italic font is sufficient to reveal dyslexics neural deficits at similar short latencies in relation to larger differences in font (i.e., Arial vs. Gigi) that evoked comparable effects in non-dyslexics, and (3) adult dyslexics likely use a word’s shape for decoding short words rapidly.

#### 4.2.2 Centrofrontal component

Following the occipitotemporal component, we identified a centrofrontal component in both of our EEG analyses starting around 210 ms post-stimulus onset. Its appearance during the analysis of both short and long italicised words indicates that it is independent of word length, and therefore, captures a more general difference associated with the processing of a variety of italicised words in dyslexia. The observed link between this component and decision performance across all participants corroborates its importance for accurate lexical decision making. In the context of our task, which explicitly asked participants to indicate whether the decision word matched its preceding sentence, it seems plausible that this component signals a combination of post-sensory processing stages such as semantic congruency and phonological awareness as part of lexical access. These processes are not mutually exclusive. They can be associated with just one time period as lexical access is inherently fast (Sereno et al., 1998) requiring the performance of crucial word identification steps within the time window of one saccade (i.e., ∼275 ms without parafoveal preview; Sereno and Rayner, 2003). Semantics, often associated with the later N400 component (Bentin et al., 1999; Holcomb and Grainger, 2006), were also found to influence lexical decision making as early as 250 ms post-stimulus (Cavalli et al., 2016).

Support for our interpretation is provided by a number of converging findings of a neural component peaking around 250 ms post-stimulus with positive topography over frontal electrodes that reflects effects of initial semantic matching of word forms or late stages of lexical access, termed recognition potential. Similar to our task, studies observing the recognition potential manipulated the congruency of terminal sentence words (Dien et al., 2003; Martín-Loeches et al., 2004) or the semantic properties of single words (Marí-Beffa et al., 2005; Martín-Loeches et al., 2001). Importantly, both the latency and amplitude of this component have been linked to reading ability on these similar tasks (Rudell and Hua, 1997).

More evidence for our centrofrontal component’s role in post-sensory processing comes from the fact that it appears independent of word length. Neural effects of word length, and therewith the physical stimulus make-up, have repeatedly been reported for an earlier time window around 100 ms post-stimulus onset (Assadollahi and Pulvermüller, 2003, 2001; Hauk et al., 2006; Hauk and Pulvermüller, 2004). In agreement with these findings, modulation of the physical stimulus properties (i.e., word length and font) were captured by our earlier occipitotemporal, but not by our centrofrontal component in dyslexia. Further support for the centrofrontal component’s role on post-sensory word identification stages comes from electrophysiological evidence of non-linguistic perceptual decision making tasks demonstrating that post-sensory neural activity is tightly linked to and a better predictor of the decision outcome than early sensory activity (Gherman and Philiastides, 2018, 2015; Philiastides et al., 2014; Philiastides and Sajda, 2006; Ratcliff et al., 2009). Congruent with the observed scalp activity profile such decision relevant post-sensory signals have been located in frontal cortices (Filimon et al., 2013; Philiastides et al., 2014, 2011). Hence, this centrofrontal component could be part of the fronto-parietal network associated with decision making.

### 4.3 Potential role of the fronto-parietal network

Both of our components overlapped in time between 209 and 236 ms post-stimulus indicating a potential functional relationship between them. In fact, the peak amplitude of the occipitotemporal component was predictive of the subsequent centrofrontal component’s peak amplitude on a single-trial basis across participants. This link illustrates that our identified components are part of a cascade of processes taking place in short succession during lexical decision making. As in previous non-linguistic decision making tasks, the information processing encoded in the early component is broadcasted onto downstream networks for subsequent post-sensory processing and decision making (Diaz et al., 2017; Philiastides and Sajda, 2006; Ratcliff et al., 2009). Attention in particular can play a crucial modulatory role during this interplay (Philiastides et al., 2006) and help facilitate the propagation and enhancement of the most diagnostic stimulus features during visual word processing (Ruz and Nobre, 2008). In this context, Amso and Scerif (2015) proposed that connections between parietal and (pre-)frontal cortex may function as continuous loops controlling executive attention and decision making whereby these loops facilitate the transformation of the early visual processing into the relevant decision evidence along the ventral stream. In our task, such top-down influence would be reflected in an enhancement, that is lower ERP amplitude, of our occipitotemporal component as shown by our control group. In this respect, dyslexics’ higher occipitotemporal ERP amplitude coupled with the centrofrontal component being linked to decision performance suggest that this network works less efficiently during word identification in adult dyslexia.

In contrast to the ERP components presented above, we did not find any significant between-group ERP differences for contrasts examining trials that presented Arial regular font. This finding suggests that, indeed, italic font affects fundamental orthographic properties of words such as word shape important during lexical decision making. If our results were reflecting general challenges of the adult dyslexic brain independent of font style, we should have observed similar neural differences for decision words presented in Arial regular font. Further, a lack of differences in later ERP components such as the P300 and N400, commonly associated with working memory and obvious semantic mismatch on linguistic tasks (e.g., Helenius et al., 1998; Van Petten, 1995), suggests that the italic font style led to the observed group-differences as opposed to different decision strategies based on other properties of our decision words (e.g. word class, expectancy or semantic incongruency). However, we cannot rule out a mediating role of these properties as we included a variety of decision words. Hence, our results show cardinally different processing in adult dyslexia occurs within 300 ms after perceiving a word.

In summary, here we contributed to the literature by revealing that even small changes in font style, as embodied by italic font, are sufficient to elicit fundamentally different neural processing within the sensory and post-sensory stages of visual word decoding in adult dyslexia. These group differences were captured by two distinct ERP components starting as early as 167 ms after the onset of a single italicised word. Here, it has become evident that font affects the rapid interplay of orthographic, lexical and semantic processes during visual word recognition, which are most likely modulated by attention. Our findings suggest refraining from using italic font in a variety of documents – especially legal contracts and education materials – in order to optimise word processing by dyslexics.

## Acknowledgements

We thank Sabina Gherman for assistance with data collection, data analysis and feedback on the manuscript. We would also like to thank Elsa Fouragnan for assistance with data analysis and Tim Reinboth for assistance with data collection.

## Declaration of interest

The authors declare no competing interests

